# From forests to fields: investigating *Culicoides* (Diptera: Ceratopogonidae) abundance and diversity in cattle pastures and adjacent woodlands

**DOI:** 10.1101/2023.08.06.552175

**Authors:** Cassandra H. Steele, Emily G. McDermott

## Abstract

*Culicoides* Latreille (Diptera: Ceratopogonidae) biting midges are hematophagous flies that feed on wild and domestic ruminants, and transmit many arboviruses, including bluetongue virus (BTV) and epizootic hemorrhagic disease virus (EHDV) circulating in the United States (US). They occupy a range of aquatic and semiaquatic habitats, and disperse short distances from their development sites. In the southeastern US there are limited studies on the abundance and diversity of *Culicoides* in wooded and adjacent livestock pasture habitats. In thus study, we characterized *Culicoides* diversity and abundance within these distinct habitat types by setting BG-Sentinel and CDC miniature suction traps baited with CO_2_ or UV-light in wooded and pasture habitats at two locations on a university beef farm in Savoy, AR. Traps were set once per week for nine weeks in August-October of 2021-2022. Fifteen species were collected during this study. The two most abundant species were *Culicoides haematopotus* Malloch and *Culicoides stellifer* Coquillett. There was a significant effect of site and location on *C. haematopotus* collections, and a significant effect and interaction of site and trap on *C. stellifer* collections. In the woods, significantly more *C. stellifer* were collected from the CDC-UV trap, while in the pasture, significantly more were collected in the CDC-CO_2_ trap. These data suggest that *C. stellifer*, a putative vector of BTV and EHDV in the southeast, may be traveling into the pasture to host-seek, while *C. haematopotus* remains primarily in the woods. This study reveals community differences between these habitat types and implications for *Culicoides* control.

## Introduction

*Culicoides* Latreille (Diptera: Ceratopogonidae) biting midges are hematophagous flies that feed on human and animal hosts (Purse et al. 2015). They are vectors of arboviruses that infect wild and domestic ruminants (Maclachlan et al 2019), including bluetongue virus (BTV) and epizootic hemorrhagic disease virus (EHDV), which are notable orbiviruses circulating in the United States (US) (Purse et al. 2015, Cottingham et al. 2021). Livestock, especially sheep, experience higher mortality from BTV infections (Maclachlan et al. 2019), and EHDV is more commonly associated with wildlife hosts, such as white-tailed deer (Stallknecht et al. 2015).

*Culicoides* that feed on multiple hosts may act as bridge vectors between livestock and wildlife communities (Talavera et al. 2018). Cattle have been implicated as reservoir hosts for BTV and EHDV (Maclachlan 1994, Rivera et al. 2021), and wild ruminants may serve as significant reservoirs for EHDV (Rivera et al. 2021). In the US, there are two confirmed vectors of BTV: *Culicoides sonorensis* Wirth & Jones and *Culicoides insignis* Lutz (Mellor et al. 2000, Pfannenstiel et al. 2015, McGregor et al. 2022). *Culicoides sonorensis* is also a confirmed EHDV vector. *Culicoides sonorensis* is primarily found in the western US in association with confinement dairy production, and *C. insignis* is a tropical species, largely confined to southern Florida (Mellor et al. 2000, Vigil et al. 2018, McGregor et al. 2022)

Bluetongue virus and EHDV transmission have not been well studied in the southeastern US, with the exception of work done on captive cervid farms in Florida (McGregor et al. 2019, Cottingham et al. 2021, McGregor et al. 2021). These studies suggest that sylvatic species, such as *Culicoides stellifer* Coquillett, may be the primary vectors in those production systems. There is less research on transmission risk to pastured livestock. An interesting complication in some parts of the southeast, including Arkansas, is the integration of forested land and cattle pastures (silvopasture), which is different than conventional captive cervid systems, which tend to be shrubbier with more plants, and less open than typical grazing pastures for livestock. It is possible that livestock raised on silvopasture, or on pastures immediately adjacent to forested land, may be at risk of BTV/EHDV transmission from sylvatic *Culicoides* vectors.

Differences in midge abundance and diversity between livestock pastures and forests have been described previously in the literature. A higher abundance of *Culicoides* have been collected in pasture habitats versus adjacent woodlands on working farms in Alabama and Brazil (Hayes et al. 1984, Carvalho et al. 2017), though collections made from a game preserve in northern Florida found similar *Culicoides* abundance on the preserve and directly adjacent natural site (McGregor et al 2021). Species composition also varies between habitat types, with species richness and diversity usually higher in the forest than adjacent pastures (Hayes et al. 1984, Carvalho et al. 2017, McGregor et al. 2021). Generally, *Culicoides* only disperse a few kilometers from their development sites in search of a mate, blood-meal, or oviposition site (Mellor et al. 2000, Kluiters et al. 2015). Female *Culicoides* may feed on a range of hosts, and some species will feed opportunistically if their preferred host is unavailable (Hopken et al. 2017, Talavera et al. 2018), which may put livestock at an increased risk if agricultural land borders wildlife areas.

The aim of this study was to examine *Culicoides* abundance within wooded habitats and adjacent pastures on a working beef farm, and to characterize community composition within each habitat type. Female *Culicoides* were tested for BTV and EHDV to begin drawing conclusions about pathogen transmission and putative vector species in the Arkansas. This study was carried out on a beef farm during the fall to coincide with peak BTV and EHDV transmission season.

## Methods

### Study sites

Two locations at the University of Arkansas Agricultural Experiment Station, Savoy Research Complex (a working beef farm in Savoy, Arkansas) were selected for this study. The Savoy Research Complex is 3,183 acres, with 2,185 acres of forest, and approximately 250 head of cattle. Both locations, hereafter referred to as Cow Calf Unit and River throughout this manuscript, were characterized by a deciduous forest and open pasture, with a distinct tree-line separating the forest and pastures (Figure 1), and were ∼ 3.38 km apart. At each location, traps were set 20 m apart at one site within the woods and at one site in the pasture. The Cow Calf Unit had a pond ∼280 m west of the woods site and held approximately 30 heifers. The River location had a pond directly adjacent to the woods site and held approximately 70 heifers. Traps were only set in pastures without cattle to prevent animals from interfering with them. Trap lines were set 43 m from the forest/pasture edge at the Cow Calf Unit, and 20 m from the edge at the River location. The two forests sites remained consistent throughout the duration of the trapping study. Due to the rotation of the herd, the pasture sites varied somewhat between weeks, but were always adjacent to the original pasture site. Traps were rotated one position down the line each week, so each trap was in each position three times, with the exception of the CDC-CO_2_ trap at the Cow Calf Unit-pasture site during week one of 2021 and the CDC-UV and CDC-CO_2_ traps during week one of 2022, which were randomly selected to exclude due to equipment issues.

**Figure 1.**
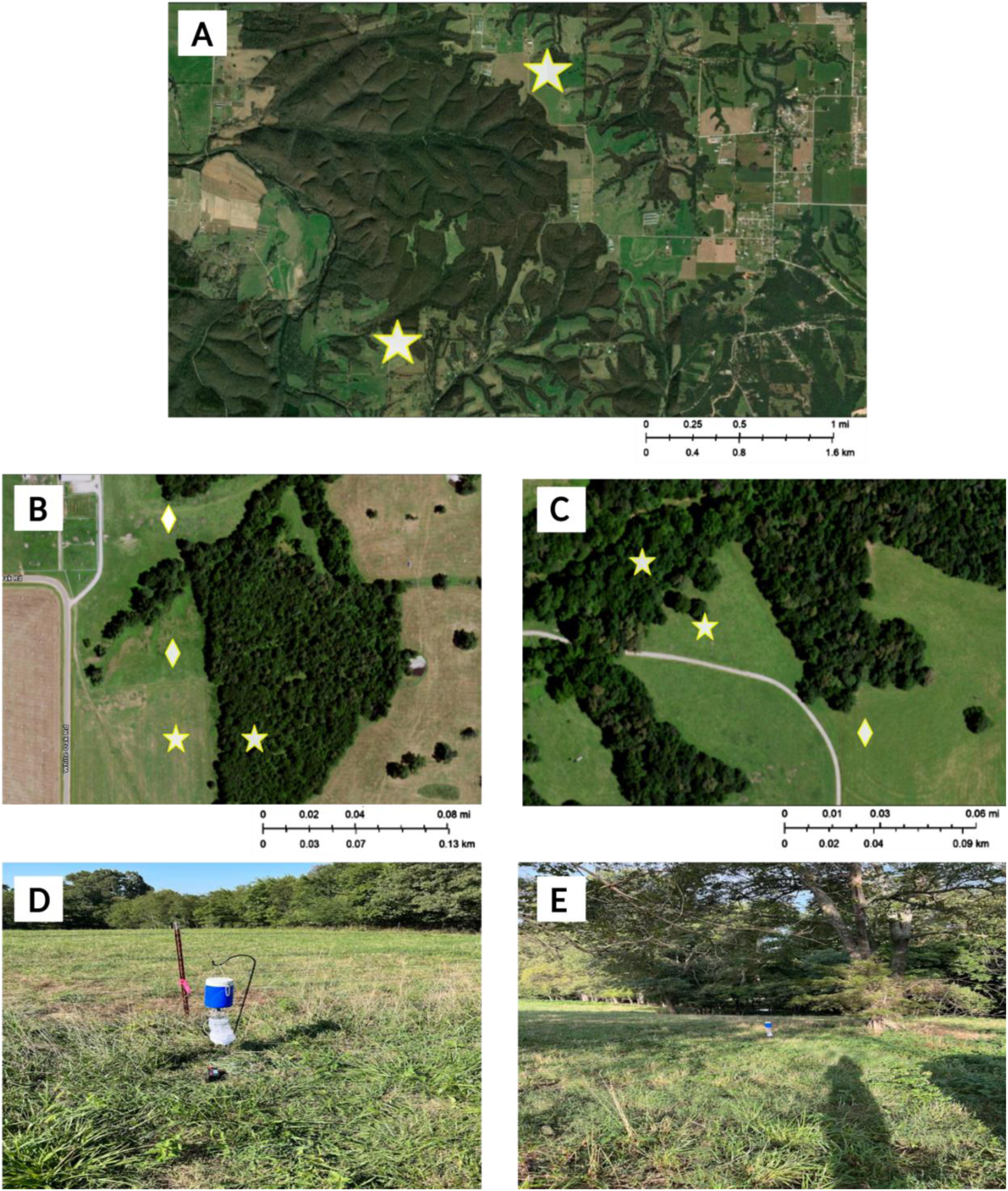
Maps of the Cow Calf Unit and River locations, represented by stars (A), as well as the pasture and woods sites within each location (B, C). In B and C, stars represent the primary trap-line, and diamonds represent alternate pasture trap-lines due to rotation of the herd. Pasture traps were set in alternative pasture during week 3 at the River location and weeks 4-5 and 7-9 at the Cow Calf Unit location in 2021, and week 1 at the Cow Calf Unit location, and weeks 5-6 at the River location in 2022. The Cow Calf Unit and River locations had a grazing pasture and integration of woodlands (D, E). Maps were created using ArcGIS Online.

### Adult <u>Culicoides</u> sampling

Trapping occurred once per week during August-October 2021 (11 August-06 October) and 2022 (22 August-19 October), for a total of nine weeks of trapping each year. In 2021, three different trap-attractant combinations were used at both the forest and pasture sites at both locations: 1) BG-Sentinel 2 (BG) trap (BioGents AG, Regensburg, Germany) baited with CO_2_, 2) CDC miniature light trap baited with UV-LED (Catalog No. 2770 BioQuip Inc., Rancho Dominguez, CA), and 3) CDC trap baited with CO_2_. Half-gallon drink coolers with a small hole drilled in the bottom were filled with ∼ 2.2 kg dry ice to disperse CO_2_. The CDC traps and dry ice cooler were suspended from shepherd’s hooks, and the CDC-CO_2_ trap was attached to the bottom of the cooler, so the release hole was just above the trap opening. The CDC-UV trap opening was ∼0.75 m from the ground, and the CDC-CO_2_ trap opening was ∼ 0.4 m from the ground. The BG trap was set directly on the ground, and the cooler was placed inside the trap on its side. The UV-LED light was disabled for the CDC-CO_2_ traps. Traps were set approximately one hour before sunset and picked up one hour after sunrise. Due to the low number of *Culicoides* collected from the BG- CO_2_ trap in 2021, this trap was excluded from year two of the study.

*Culicoides* were collected into dry trap bags and transported back to the lab where they were immediately placed in the freezer for ∼1 h to kill the specimens before sorting. Collections were stored dry at -20°C for 1-2 d, at which time *Culicoides* were removed, transferred to 70% ethanol, and stored at -20°C until identification. *Culicoides* were identified based on wing pattern using the keys of Jamnback (1965), Battle & Turner (1971), Wirth & Blanton (1987), and Wirth et al. (1985), and sorted by sex and parity (Akey & Potter 1979). Identified midges were stored in RNA*later* stabilization solution (Thermo Fisher Scientific, Waltham, MA) at -20°C until processing for BTV/EHDV detection.

### BTV/EHDV detection

Parous female *Culicoides* were stored in pools of up to 20 individuals. Pools were separated based on collection date, location, site, trap, and species. Midges were homogenized using the Next Advance Navy bead lysis kit (Next Advance, Troy, NY) with 500 μl of lysis/binding concentrate buffer solution (Thermo Fisher Scientific, Waltham, MA) in a Bullet Blender STORM (Next Advance Inc., Troy, NY). Samples were homogenized for 3 min at speed 12. Viral RNA was extracted from homogenized samples using the MagMax -96 Viral RNA Isolation kit (Thermo Fisher Scientific, Waltham, MA). Homogenized samples were stored at - 80°C.

Pools were tested for BTV and EHDV using a previously published multiplex qRT-PCR assay (Wernicke et al. 2015, McGregor et al. 2019). 5 μL of template RNA was combined with 5 μL TaqMan Fast Virus One Step Master Mix (Thermo Fisher Scientific, Waltham, MA), 4.55 μL molecular grade water, 0.5 μM each BTV forward and reverse primers, 0.75 μM each EHDV forward and reverse primers, 0.1 μM FAM-labelled BTV probe, and 0.125 μM VIC-labelled EHDV probe in a 20 μL reaction. Reactions were run using a QuantStudio^TM^ 3 Real-Time PCR System (Applied Biosystems^TM^, Waltham, MA) with the following cycling conditions: 50° C for 5 min, 95° C for 20 sec, followed by 40 cycles of 95° C for 3 sec, and 60° C for 30 sec. Extracted RNA from stocks of BTV-10 (National Veterinary Services Laboratories 001-ODV) and EHDV- 1 (National Veterinary Services Laboratory 040-ODV) were included in every plate as a positive controls, and ddH_2_O was included as a negative control. Ct values of <31 were considered positive.

### Statistical analysis

All analyses were conducted using R (v. 4.2.3). A generalized linear model (GLM) with a negative binomial distribution (package “MASS”, Ripley et al. 2023, R package version 7.3-60) was used to compare the effect and interactions of year, site, trap, and location on females (pars and nullipars combined) abundance of the two most frequently collected species, *Culicoides haematoptous* Malloch and *C. stellifer.* Since these species comprised the majority of the collections, effects of year, site, trap, and location on total *Culicoides*, or on other individual species, were not analyzed statistically. All factors were included as fixed effects and the BG-CO_2_ trap data were dropped from the analysis due to the negligible number of *Culicoides* collected. Initial models included all interaction terms, which were dropped in the order of decreasing significance (P>0.05). Model fit was assessed using an ANOVA Chi-square test and comparison of AIC scores. If there was no significant difference between models (P>0.05), the simplest model with the lowest AIC was chosen. For *C. stellifer* the final model included location, site, trap, and their interactions (AIC: 231), and for *C. haematopotus*, the final model included year, site, trap, and location (AIC: 296). To examine the significant interaction between site and trap on *C. stellifer* abundance, the emmeans function was used to test multiple comparisons (package “emmeans”, Russell et al. 2023, R package version 1.8.6). *Culicoides stellifer* abundance in the woods versus the pasture across both locations and years was also analyzed using a negative binomial distribution model to compare the effect of trap on both sites separately.

The weekly Shannon diversity index (H) was calculated using the sum of collections from years one and two for Cow Calf Unit-pasture, Cow Calf Unit-woods, River-pasture, and River-woods using the package ‘vegan’ (Okansen et al. 2022, R package version 2.6-4). A Kruskal-Wallis rank sum test was used to compare the differences in the mean weekly Shannon index, followed by a Dunn test (package ‘dunn.test’; Dinno 2017, R package version 1.3.5) for means separation. The overall Shannon index for each location and site across both years was calculated as well. To determine the differences between communities at each location and site, the Morisita-Horn similarity index (C_H_) was calculated using package ‘divo’ (Sadee et al. 2016, R package version 1.0.1).

## Results

### <u>Culicoides</u> Abundance

In total, 989 *Culicoides* representing 15 species were collected across both years of the study; (year one: n= 584, year two: n= 405). *Culicoides haematopotus* (total: n = 753) and *C. stellifer* (total: n = 138) made up a majority of the *Culicoides* collected, however there were a number of other species collected, including *Culicoides crepuscularis* Malloch (n = 40), *Culicoides variipennis* Coquillett (n = 24), and *Culicoides venustus* Fabricius (n = 15) (Table 1). The remaining species were primarily collected from the River woods site, and comprised 11% of the total collection.

**Table 1.** Total male and female *Culicoides* sampled from August-October 2021 and 2022.

| Species | 2021 | 2022 |
| --- | --- | --- |
| *C. arboricola* | 1 | 6 |
| *C. baueri*<sup>1</sup> | 4 | 3 |
| *C. bickleyi* | 2 | 0 |
| *C. crepuscularis* | 15 | 25 |
| *C. debilipalpis*<sup>1</sup> | 1 | 0 |
| *C. guttipennis* | 2 | 3 |
| *C. haematopotus* | 485 | 258 |
| *C. hinmani*<sup>2</sup> | 1 | 0 |
| *C. nanus*<sup>1</sup> | 1 | 0 |
| *C. ousairani*<sup>1</sup> | 0 | 1 |
| *C. obsoletus* group | 1 | 0 |
| *C. oklahomensis*<sup>1</sup> | 3 | 0 |
| *C. stellifer* | 70 | 68 |
| *C. variipennis* | 4 | 20 |
| *C. venustul*<sup>1</sup> | 11 | 4 |
| **Total** | **601** | **388** |

<sup>1</sup>Species only collected from River location
<sup>2</sup>Species only collected from Cow Calf Unit

Collections with the highest average diversity were made from Cow Calf Unit-pasture (H=0.63±0.31), River-pasture (H=0.69±0.57), and River-woods (H=0.63±0.32), with Cow Calf Unit-woods having the lowest average diversity (H=0.19±0.34) (Table 2).

**Table 2.** Mean Shannon diversity index (H) of site and location in year one and two. Standard deviation (SD) in parentheses. Superscript letters denote significant differences between location-site combinations (P<0.05).

| Location | Site | Mean Shannon Index (SD) |
| --- | --- | --- |
| Cow Calf Unit | Pasture | 0.626±0.31<sup>a</sup> |
|  | Woods | 0.192±0.34<sup>b</sup> |
| River | Pasture | 0.695±0.57<sup>a</sup> |
|  | Woods | 0.633±0.32<sup>a</sup> |

The Cow Calf Unit-woods site was significantly less diverse (P≤0.04) than the remaining locations and sites, which were not significantly different from each other. The Morisita-Horn similarity index revealed that the two woods sites and the two pasture sites were composed of very similar species communities (Cow Calf Unit-woods/River-woods: C_H_ = 0.99, Cow Calf Unit-pasture/River-pasture: C_H_ = 0.70). The Cow Calf Unit-woods and River-pasture sites had the least overlap between species communities (C_H_ = 0.36) (Table 3).

**Table 3.** The Morisita-Horn similarity index (C_H_) of location/site diversity comparisons across both years of this study. 95% confidence intervals are presented in parentheses. C_H_ ranges from 0 (no overlap between communities) to 1 (complete overlap between communities).

|  | River/Pasture | River/Woods | CCU/Pasture |
| --- | --- | --- | --- |
| River/Woods | 0.426 (0.28,0.59) |  |  |
| CCU/Pasture | 0.703 (0.57,0.86) | 0.465 (0.41,0.48) |  |
| CCU/Woods | 0.36 (0.20,0.52) | 0.992 (0.991,0.992) | 0.41 (0.35,0.42) |

Averaging both years of the study, the River location collected a higher mean abundance of total midges per trap night (n = 5.12 ± 18.7 [SD]) than the Cow Calf Unit (1.20 ± 3.14) (Supplemental file 1). Additionally, woods sites had a higher abundance (5.36 ± 19.1) than the pasture sites (1.02 ± 1.59) (Table 4). The CDC-UV trap collected a higher mean abundance of midges (5.83 ± 19.4) than the BG-CO_2_ (0.31 ± 0.58) and CDC-CO_2_ traps (0.76 ± 1.72).

**Table 4.** Total abundance of male and female *Culicoides* sampled per trap night from the Cow Calf Unit and River locations in August-October 2021 and 2022. Average abundance (±SD) per trap night is presented in parentheses.

| Species | Pasture Collections | Woods Collections |
| --- | --- | --- |
| *C. arboricola* | 0 | 7 (0.043 ± 0.26) |
| *C. baueri* | 1 (0.009±0.094) | 6 (0.037±0.22) |
| *C. bickleyi* | 1 (0.009±0.094) | 1 (0.006±0.079) |
| *C. crepuscularis* | 16 (0.14±0.37) | 24 (0.15±0.76) |
| *C. debilipalpis* | 1 (0.009±0.094) | 0 |
| *C. guttipennis* | 1 (0.009±0.094) | 4 (0.025±0.16) |
| *C. haematopotus* | 24 (0.21±0.78) | 719 (4.41±19.2) |
| *C. hinmani* | 0 | 1 (0.006±0.079) |
| *C. nanus* | 0 | 1 (0.006±0.079) |
| *C. obsoletus group* | 0 | 1 (0.006±0.079) |
| *C. oklahomensis* | 0 | 3 (0.018±0.13) |
| *C. ousairani* | 0 | 1 (0.006±0.079) |
| *C. stellifer* | 47 (0.42±1.13) | 91 (0.56±2.57) |
| *C. variipennis* | 19 (0.17±1.08) | 5 (0.031±0.32) |
| *C. venustus* | 5 (0.04±0.25) | 10 (0.06±0.45) |
| **Total** | **115** | **874** |

*Culicoides haematopotus* and *C. stellifer* were the most abundant species, representing 75.1 and 13.9% of the total collections, respectively. There were no significant interactions between site, trap, location, and year on *C. haematopotus* abundance, so interaction terms were dropped. There was no significant difference in female *C. haematopotu*s abundance between 2021 and 2022, so data from both years of the study were combined for analysis. There was no significant difference in the number of female *C. haematopotus* collected in CDC traps baited with CO_2_ or UV. There was a significant effect of site and location on *C. haematopotus* abundance, with more being collected on average from the River location per trap night (9.39 ± 20.7) than from the Cow Calf Unit (0.80 ± 3.72; P=0.005) (Figure 2A), and more collected in the woods sites per trap night (9.61 ± 30.3) than the pasture sites (0.32 ± 0.98; P<0.001) (Figure 2B).

**Figure 2.**
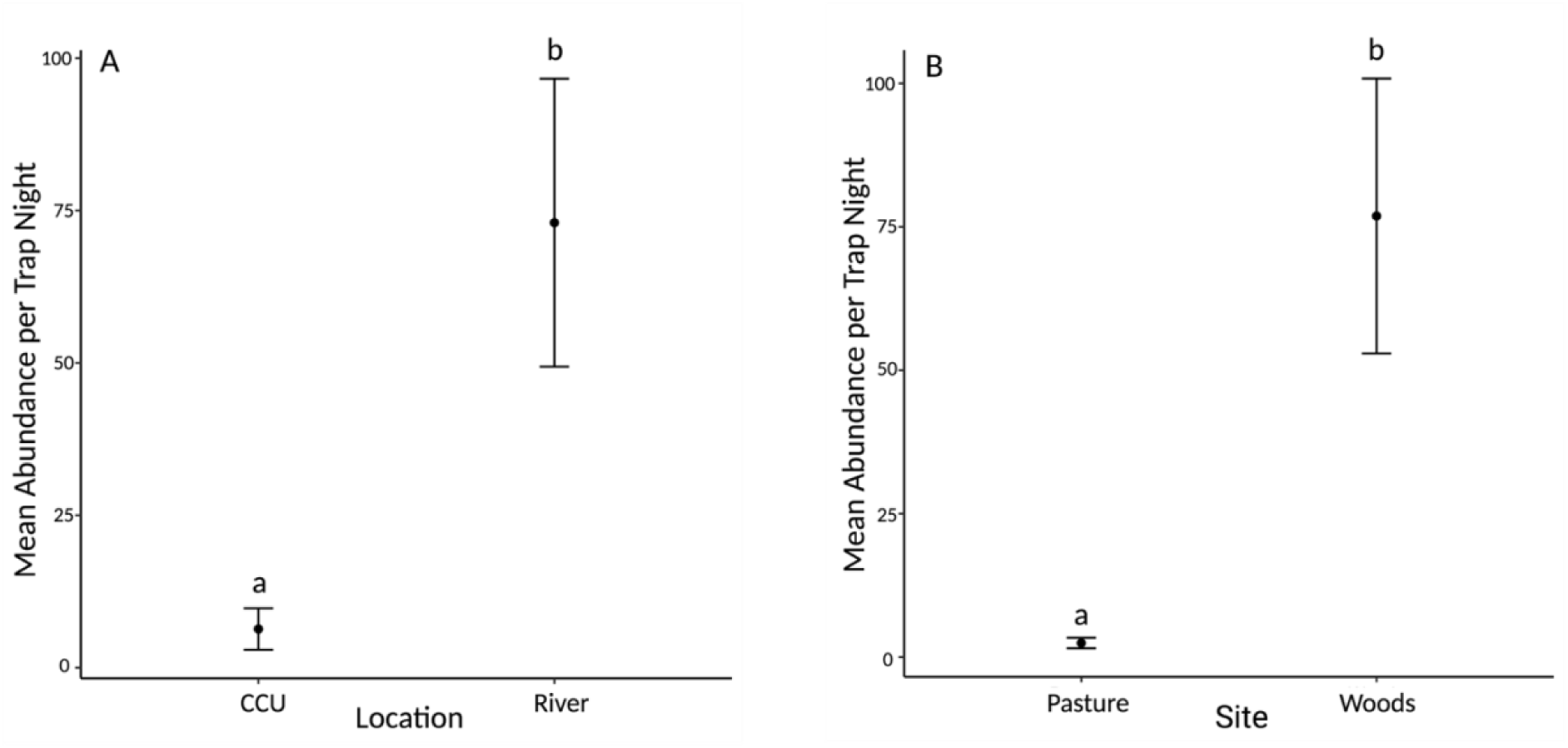
Mean abundance (±SD) per trap night of total female *C. haematopotus* collected from the Cow Calf Unit and River locations (A) and from the pastures vs woods (B). Letters represent significant differences between locations or sites (P<0.05).

For female *C. stellifer*, there was no significant effect of year or location on abundance, so data were averaged across 2021 and 2022, and across River and Cow Calf Unit locations for analyses. Trap and site did significantly affect the mean abundance of *C. stellifer* collected (P≤0.01). Significantly more midges were collected on average in woods sites per trap night (1.20 ± 4.55) than in pasture sites (0.61 ± 1.92; P=0.08), and significantly more were collected in CDC-UV traps (1.27 ± 4.58) than CDC-CO_2_ traps (0.54 ± 1.90; P=0.01). There was also a significant interaction between site and trap, such that in woods sites, more female *C. stellifer* were collected in the UV traps than CO_2_ traps on average per trap night (2.33 ± 6.27; P<0.001), but in pasture sites, significantly more were collected in the CO_2_ traps (1.06 ± 2.64; P=0.012) (Figure 3).

**Figure 3.**
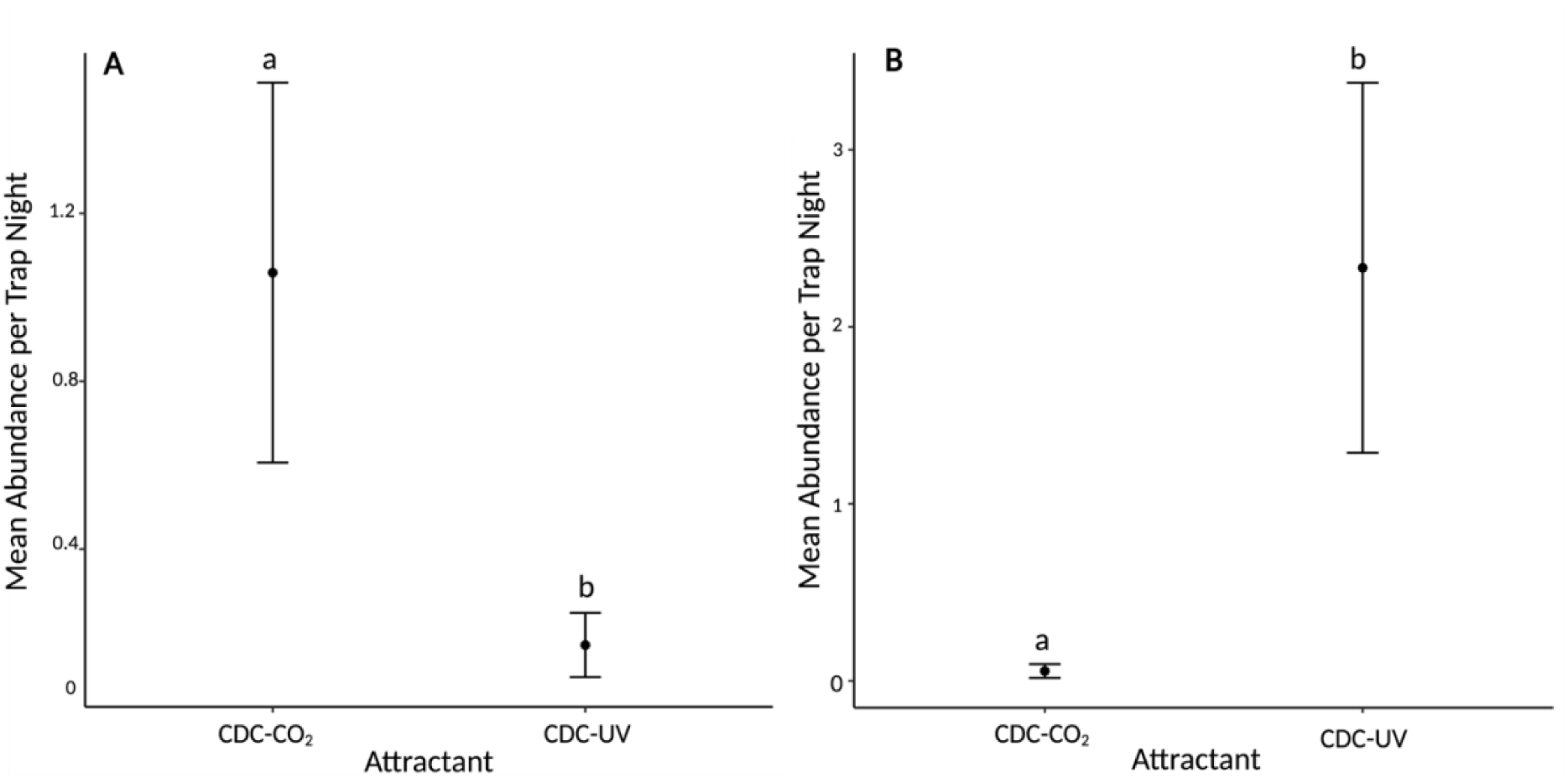
Mean abundance (±SD) per trap night of *C. stellifer* collected from the pasture (A) and woods (B) in the CDC-CO_2_ versus CDC-UV traps. Letters represent significant differences between attractants (P<0.05).

### Pathogen Detection

117 pools of parous females representing 13 species from 2021-2022 were tested for BTV and EHDV. No pools were positive for BTV or EHDV (Supplemental File 2).

## Discussion

Many *Culicoides* readily feed on, and transmit pathogens to, cattle, but the majority of work on BTV transmission has focused on confinement, or feedlot, dairy production. There is limited information about the *Culicoides* species present in grazing pastures for cattle, or in forested habitats adjacent to those pastures. To answer the question of whether cattle in cow-calf production systems in northwest Arkansas are at risk of BTV/EHDV transmission, we collected *Culicoides* from these distinct habitat types on a working beef farm. The findings of this study suggest that *C. stellifer*, a putative vector of hemorrhagic disease viruses (McGregor et al. 2019), is present in the woods and in the pasture, and that *C. stellifer* may be dispersing out of the woods to host-seek. *Culicoides haematopotus*, the most abundant species collected, was found primarily in the woods, suggesting they are likely not dispersing out into the pasture to host-seek and may not have a major role in pathogen transmission.

Other studies have characterized the diversity and abundance of *Culicoides* spp. in livestock pastures and wooded habitats. Many of these studies describe higher diversity in woods collections, but the effect of agricultural versus natural habitat on *Culicoides* abundance is less clear. Species richness was found to be higher in forest habitats in a study from Brazil, but a greater number of *Culicoides* were collected from an adjacent pasture (Carvalho et al. 2017). Rigot et al. (2013) examined Paleartic *Culicoides* abundance on dairy farms and the surrounding landscape and found that abundance decreased as distance from the farm increased. However, abundance increased again in the woodland habitat, indicating that *Culicoides* prefer to stay near potential hosts, whether they are domesticated or wildlife. In Florida, McGregor et al. (2021) found that *Culicoides* collections were the most similar between a big game preserve with various species of Bovidae and Cervidae and a directly adjacent site within a state forest bordering the preserve, with abundance decreasing away from the preserve. Additionally, for *C. stellifer*, there was a greater abundance of parous and blood fed midges collected in the preserve and adjacent site, suggesting that female midges may move into the preserve in search of a host. Species richness was also greater in natural sites than in the preserve (McGregor et al. 2021).

Our study supports the idea that wooded areas support a greater diversity of *Culicoides*. We found high diversity in the woods, with 14/15 species collected, compared to 9/15 species collected in the pastures, and overall a high abundance from the woods (n = 874) and low abundance in the pastures (n = 115). In both habitat types, *C. haematoptous* and *C. stellifer* were the most abundant species collected.

The River-pasture, River-woods, and Cow Calf Unit-pasture sites had the highest diversity of *Culicoides*, and each site had similar mean Shannon diversity indices. The River sites were much wetter than the Cow Calf Unit sites, with a small pond and stream directly adjacent to the woods and pasture. The majority of *Culicoides* collected during the study also came from the River sites. The higher diversity of midges collected from the Cow Calf Unit- pasture versus the woods site is surprising. The Cow Calf Unit-pasture was much drier, and there was less vegetation and shade in the woods than at the River sites. While there were no obvious development sites in the pasture or woods at the Cow Calf Unit, it is possible that midges were developing somewhere nearby and traveling into the pasture in search of a host. The species communities were similar between the two pasture and two woods sites, with there being the least species overlap between the Cow Calf Unit-woods and River-pasture site. These two habitats were contrastive, with the River-pasture having taller grass and some shade, with an adjacent pond, while the Cow Calf Unit-woods was dry, with tall trees and less ground vegetation. The Cow Calf Unit-woods and River-woods had high overlap between communities as well. Both sites were shaded, however the River-woods site was wetter and more vegetated than the Cow Calf Unit-woods, and the highest abundance of midges were collected from the River-woods site.

*Culicoides haematopotus* was primarily collected from the woods, but was also the second most common species collected in the pastures. Although this species is thought to primarily feed on birds, *C. haematoptous* has been collected in association with large mammals, including deer (Smith & Stallknecht 1996, Swanson & Turnbull 2014), and it will feed on deer, as well as birds (McGregor et al. 2019). Perhaps due to this opportunistic feeding behavior, BTV has been detected from pools of field-collected *C. haematopotus* in Louisiana (Becker et al. 2010), but its role in potential BTV/EHDV transmission is unclear (McGregor et al. 2022). The abundance of *C. haematopotus* collected in the woods in this study reinforces that this species is likely ornithophilic and will remain in wooded areas where their preferred hosts are present.

However, *C. haematopotus* presence in the pastures (20% of pasture collections) suggests they may be more opportunistic than we believe. Wood-nesting birds are unlikely to nest or be found in a pasture habitat around dawn and dusk (Wegner & Merriam 1979), which are peak *Culicoides* activity hours (Mellor et al. 2000), however field-nesting species or livestock and wildlife present in the pasture may encourage *C. haematopotus* to travel out of the woods (Wegner & Merriam 1979).

*Culicoides stellifer*, the second most abundant species collected in our study, is a putative vector of BTV and EHDV in the southeastern US (McGregor et al. 2019). *Culicoides stellifer* is a sylvatic species that is widely distributed across the US, and readily feeds on ungulate hosts (McGregor et al. 2022). A study in Alabama found a higher abundance of *C. stellifer* collected off cattle in a pasture than in adjacent woodlands (Hayes et al. 1984). We found that there were more *C. stellifer* collected from the woods using the UV attractant and more collected from the pasture when traps were baited with CO_2_ (Figure 4), suggesting that the midges are developing/resting in the woods and host-seeking in the pasture. It is unlikely that *C. stellifer* were developing in the pastures where we conducted our study due to the unavailability of suitable larval development sites. *Culicoides stellifer* is often associated with shaded development sites with organic debris and leaf litter (Erram et al. 2019). The pastures at both the River and Cow Calf Unit locations were extremely dry during August-October in both years of this study. Common potential development sites, such as mud under water troughs or ponds, were not present in the immediate vicinity of the pasture traps at the Cow-Calf Unit. A small, shaded pond was present directly adjacent to the woods and pasture sites at River, and emergence samples collected from this pond at the same time as this study did yield a small number of adult *C. stellifer*. A pond located ∼200 m from the Cow Calf Unit location was also sampled but did not produce any adult midges. Woods sites at both locations were also usually dry, particularly at the Cow Calf Unit location, but were not exhaustively searched for potential development sites, and so suitable, cryptic sites may have been present.

*Culicoides stellifer* is a likely vector for BTV/EHDV transmission. There have been positive pools of BTV and EHDV from field-collected adults in Florida (McGregor et al. 2022) and Louisiana (Becker et al. 2020), and this species satisfies two of the World Health Organization vector incrimination criteria in North America (McGregor et al. 2022).

Additionally, unlike *C. haematopotus*, *C. stellifer* has been collected from light traps near deer enclosures and directly off the deer themselves (Smith & Stallknecht 1996), and frequently feeds on susceptible ruminants (McGregor et al. 2018). This provides further evidence that *C. stellifer* may be moving out of the woods to find a host and potentially transmitting pathogens (Hopken et al. 2017, McGregor et al. 2019).

Other *Culicoides* species that were collected in this study may play a role in virus transmission. *Culicoides crepuscularis* (4% total collection) is considered to be ornithophilic, however BTV has been detected in field-collected females in Louisiana (Becker et al. 2014), and this species has been reported to feed on ungulate hosts, such as white-tailed deer (Smith & Stallknecht 1996, Hopken et al. 2017). *Culicoides variipennis* is closely related to *C. sonorensis*, a confirmed vector of BTV and EHDV, however the role of *C. variipennis* in transmission is unclear (McGregor et al. 2022). Tabachnick (1996) reported low BTV infection rates in field-collected *C. variipennis* in the northeast US, but pooled midges from Louisiana were negative for both BTV and EHDV (Becker et al. 2020). We did not detect either virus in *C. variipennis* but did collect this species in a higher proportional abundance in the pastures (2% of pasture collections) than the woods (0.5% of woods collections), suggesting it feeds preferentially on susceptible livestock, in line with what is known about this species’ host preference (Hopken et al. 2017). Field data have implicated *C. venustus* as a potential vector of EHDV. Field-collected *C. venustus* have tested positive for EHDV in Florida (McGregor et al. 2019) and BTV in Louisiana (Becker et al. 2021), and this species does feed on white-tailed deer (McGregor et al. 2019). We collected *C. venustus* from the pasture and the woods (1.5% of total collections), suggesting that this species may disperse out of the woods to locate a suitable host. However, *C. venustus* was found in a low abundance in this study (n = 15), compared to McGregor et al. (2019) that collected more in Florida (n = 873). *C. venustus* role in pathogen transmission is not well understood, but they may be more important vectors in areas that have a higher abundance, such as in Florida.

Typical cultural control of *Culicoides* consists of eliminating development sites and stabling animals (Mullens et al. 2015). Previous studies have attempted to eliminate development sites but saw little reduction in *Culicoides* abundance, likely due to immigration from neighboring farms, other breeding sites, or adjacent wooded areas (Mellor et al. 2000, Mayo et al. 2014, Harrup et al. 2016). Separating livestock from *Culicoides* is important to reduce virus transmission, however there is limited information on the feeding habits of sylvatic species and whether or not they will readily enter stables to feed (Pfannenstiel et al. 2015). Our results suggest that some *Culicoides* species are moving out of the woods to blood-feed, especially since the pastures have no obvious available oviposition sites. *Culicoides* usually do not travel far to host-seek (Mellor et al. 2000), however, it has been shown that *C. sonorensis* can disperse at least 2.8 km in one night (Pfannenstiel et al. 2015), for example. The dispersal of important sylvatic species, such as *C. stellifer*, into open fields and pastures is poorly understood, but mosquito behavior may provide insight into this phenomenon.

Mosquitoes may use the woodland edge as a visual cue to move in and out of a forested habitat. Some species remain in the forest during the day but disperse into an open field at night (Bidlingmayer & Hem 1981). Mosquitoes’, and other vectors’, distribution across an open field, forest/field edge, and forest may depend on host density and preference, larval habitat availability, and plant nectar sources (Reiskind et al. 2017). In this study, *C. haematopotus* dominated the woods collections, and this species is usually associated with birds (Hopken et al. 2017). There were no obvious larval habitats in the open pasture, so sylvatic *Culicoides*, such as *C. stellifer*, that were present in the pasture, may be using the woods/pasture edge as a visual cue to move out to find a blood meal. In our study, *C. stellifer* were predominantly collected from the pasture in CDC-CO_2_ traps, indicating that they are host-seeking, and collected from the woods in CDC-UV traps, where they are likely resting and ovipositing. Other species, such as *Culicoides guttipennis* Coquillett, that develop in tree holes, were predominantly found in the woods and not in the pasture, indicating that there may be some habitat fidelity among *Culicoides* species. Lastly, although the vegetation and available sugar sources at our study locations were not recorded, the pasture was predominately grass with few flowering plants, so it is unlikely that midges were traveling out of the woods to find a nectar source.

Some mosquito species are also influenced by a phenomenon known as edge effects, wherein some species are more common in this transition region between different biological communities (Reiskind et al. 2017). It is possible that *Culicoides* may exhibit similar behavior, since some species, such as *Culicoides arboricola* Root & Hoffman and *C. guttipennis*, were almost strictly found in the woods, while other species, such as *C. stellifer* and *C. venustus*, were found in the woods and pasture habitats. The community found at the forest/pasture edge was not examined in this study, but future work will involve setting traps in this ecotone to determine if there is an effect on collected species. The effect of the woods/pasture edge on *Culicoides* movement between habitat types is not well understood, but it may be beneficial for midge control to move fence lines and keep livestock away from the woodlands, which may aid in reduced biting risk and subsequent pathogen transmission.

Seasonal pasture rotation may also aid in reducing biting risk by limiting *Culicoides* host contact. Pasture rotation can be used as an internal parasite control for grazing livestock (Barger 1999, Relf et al. 2012), so similar methods could be implemented for midge control. *Culicoides* bite throughout the warm months, however peak BTV and EHDV transmission is in the fall (Pfannenstiel et al. 2015), so rotating livestock away from the woods during this period of time may aid in reducing virus transmission. Future work will also include determining if *Culicoides* are moving out of the woods to host seek and how far they are actually traveling.

This study provides a detailed overview of *Culicoides* species composition in pastures and adjacent woodlands on a working beef farm. *Culicoides haematopotus* and *C. stellifer* were the two most abundant species, and based on the differences in *C. stellifer* collections in UV and CO_2_ traps set in the woods versus the pastures, it is likely this species is moving between these habitat types to locate hosts. There were no positive detections of BTV/EHDV during this study, likely due to the low number of total *Culicoides* collected. However, our results demonstrate community differences between pasture and wooded habitats and the potential implications this has for control of *Culicoides*.

## Acknowledgments

We thank the University of Arkansas System Division of Agriculture and the Department of Entomology and Plant Pathology for their continued support in this project, the University of Arkansas Savoy Research Center for permission to trap, Bernard Krumpelman for assisting us with scouting trap locations, Cierra Briggs for providing trapping materials and assistance, as well as the McDermott lab for their continued support and help throughout this project. This work was supported by USDA-NIFA Hatch Project ARK02726. Figures were created using BioRender.com.

## SUPPLEMENTARY MATERIAL

**Supplemental File 1.** Total abundance of male and female *Culicoides* collected from Cow Calf Unit and River locations in 2021-2022.

| Species | Cow Calf Unit | River |
| --- | --- | --- |
| *C. arboricola* | 1 | 6 |
| *C. baueri* | 0 | 7 |
| *C. bickleyi* | 1 | 1 |
| *C. crepuscularis* | 10 | 30 |
| *C. debilipalpis* | 0 | 1 |
| *C. guttipennis* | 1 | 4 |
| *C. haematopotus* | 72 | 671 |
| *C. hinmani* | 1 | 0 |
| *C. nanus* | 0 | 1 |
| *C. obsoletus group* | 0 | 1 |
| *C. ousairani* | 0 | 1 |
| *C. oklahomensis* | 0 | 3 |
| *C. stellifer* | 18 | 120 |
| *C. variipennis* | 16 | 8 |
| *C. venustus* | 0 | 15 |
| **Total** | **120** | **869** |

**Supplemental file 2.**
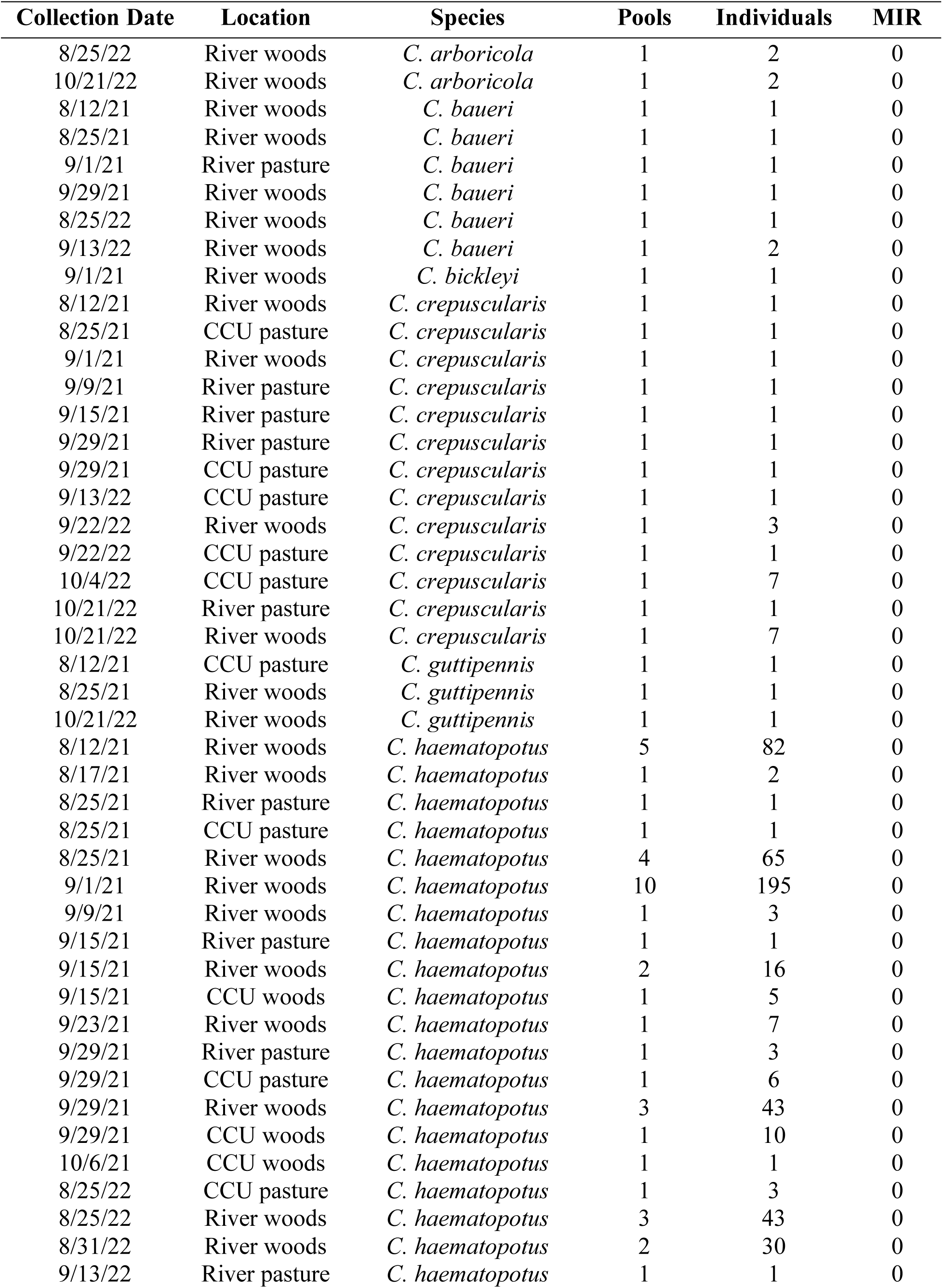
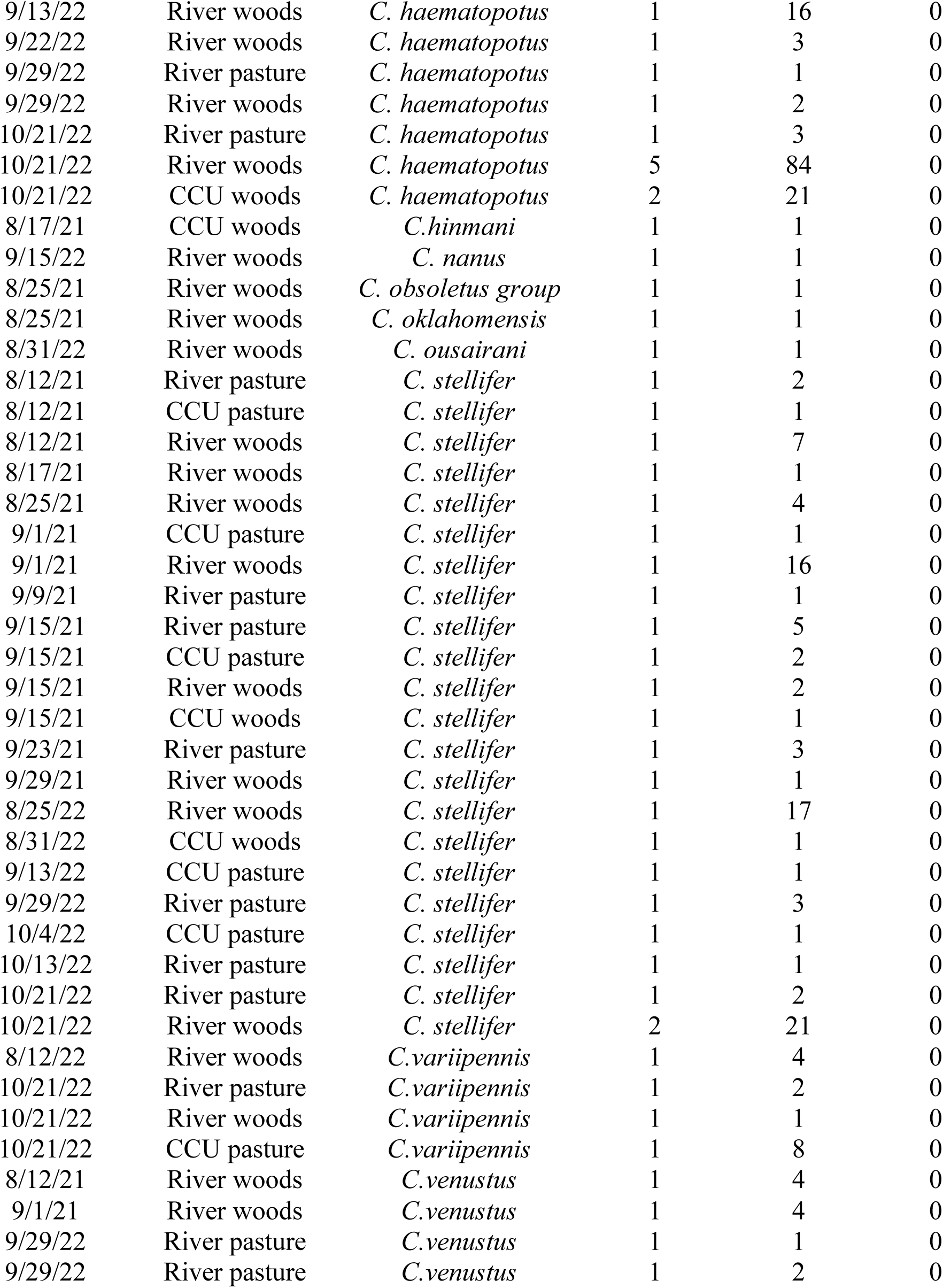
Minimum infection rate of parous *Culicoides* females tested for BTV and EHDV. CCU: Cow Calf Unit.

## References

Barger, I. A. 1999. The role of epidemiological knowledge and grazing management for helminth control in small ruminants. International journal for parasitology, 29(1): 41–47.

Battle, W. R., and E.C. Turner. 1971. A systematic review of the genus *Culicoides*. The Insects of Virginia: no. 3. 1–129.

Becker, M. E., W.K. Reeves, S.K. Dejean, M.P. Emery, E.N. Ostlund, and L.D. Foil. 2014. Detection of bluetongue virus RNA in field-collected *Culicoides* spp.(Diptera: Ceratopogonidae) following the discovery of bluetongue virus serotype 1 in white-tailed deer and cattle in Louisiana. Journal of medical entomology. 47(2): 269–273.

Becker, M. E., J. Roberts, M.E. Schroeder, G. Gentry, and L.D. Foil. 2020. Prospective study of epizootic hemorrhagic disease virus and bluetongue virus transmission in captive ruminants. Journal of medical entomology. 57(4): 1277–1285.

Becker, M., J.S. Park, G. Gentry, C. Husseneder, and L. Foil. 2021. Comparison of trapping methods for use in surveys for potential *Culicoides* vectors of orbiviruses. Parasites & Vectors. 14: 1–11.

Bidlingmayer, W. L., and D.G. Hem. 1981. Mosquito flight paths in relation to the environment: effect of the forest edge upon trap catches in the field. Mosquito News. 41: 55–59

Carvalho, L.P.C., A.M. Pereira Júnior, E.D.S. Farias, J.F. Almeida, M.S. Rodrigues, F. Resadore, F.A.C. Pessoa, and J.F.D. Medeiros. 2017. A study of *Culicoides* in Rondônia, in the Brazilian Amazon: species composition, relative abundance and potential vectors. Medical and Veterinary Entomology. 31(1): 117–122.

Cottingham, S. L., Z.S. White, S.M. Wisely, and J.M. Campos-Krauer. 2021. A Mortality-based description of EHDV and BTV prevalence in farmed white-tailed deer (Odocoileus virginianus) in Florida, USA. Viruses. 13(8): 1443.

Erram, D., E.M. Blosser, N. Burkett-Cadena. 2019. Habitat associations of *Culicoides* species (Diptera: Ceratopogonidae) abundant on a commercial cervid farm in Florida, USA. Parasites & vectors. 12: 1–13.

Harrup, L. E., M.A. Miranda, and S. Carpenter, S. 2016. Advances in control techniques for *Culicoides* and future prospects. Vet Ital. 52(3-4): 247–64.

Hayes, M.E., G.R. Mullen, and K.E. Nusbaum. 1984. Comparison of *Culicoides* spp.(Diptera: Ceratopogonidae) attracted to cattle in an open pasture and bordering woodland. Mosquito news (USA). 44:368–370.

Hopken, M.W., B.M. Ryan, K.P. Huyvaert, and A.J. Piaggio. 2017. Picky eaters are rare: DNA- based blood meal analysis of *Culicoides* (Diptera: Ceratopogonidae) species from the United States. Parasites & Vectors. 10:1–9.

Jamnback, H. 1966. The Culicoides of New York State (Diptera: Ceratopogonidae). Bulletin New York State Museum and Science Service. 399:1–154.

Kluiters, G., H. Swales, H., and M. Baylis. 2015. Local dispersal of palaearctic *Culicoides* biting midges estimated by mark-release-recapture. Parasites & vectors. 8(1): 1–9.

MacLachlan, N.J. 1994. The pathogenesis and immunology of bluetongue virus infection of ruminants. Comparative immunology, microbiology and infectious diseases. 17(3-4): 197–206.

Maclachlan, N.J., S. Zientara, W.C. Wilson, J.A. Richt, and G. Savini. 2019. Bluetongue and epizootic hemorrhagic disease viruses: recent developments with these globally re-emerging arboviral infections of ruminants. Current Opinion in Virology. 34: 56–62.

Mellor, P.S., J. Boorman, and M. Baylis. 2000. *Culicoides* biting midges: their role as arbovirus vectors. Annual review of entomology. 45(1): 307–340.

McGregor, B.L., K.E. Sloyer, K.A. Sayler, O. Goodfriend, J.M.C. Krauer, C. Acevedo, X. Zhang, D. Mathias, S.M. Wisely, and N.D. Burkett-Cadena. 2019. Field data implicating *Culicoides stellifer* and *Culicoides venustus* (Diptera: Ceratopogonidae) as vectors of epizootic hemorrhagic disease virus. Parasites & vectors. 12: 1–13.

McGregor, B. L., T. Stenn, K.A. Sayler, E.M. Blosser, J.K Blackburn, S.M. Wisely, and N.D. Burkett-Cadena. 2019. Host use patterns of *Culicoides* spp. biting midges at a big game preserve in Florida, USA, and implications for the transmission of orbiviruses. Medical and veterinary entomology. 33(1): 110–120.

McGregor, B.L., J.K. Blackburn, S.M. Wisely, and N.D. Burkett-Cadena. 2021. *Culicoides* (Diptera: Ceratopogonidae) communities differ between a game preserve and nearby natural areas in northern Florida. Journal of Medical Entomology. 58(1): 450–457.

McGregor, B. L., P.T. Shults, and E.G. McDermott. 2022. A review of the vector status of North American *Culicoides* (Diptera: Ceratopogonidae) for bluetongue virus, epizootic hemorrhagic disease virus, and other arboviruses of concern. Current Tropical Medicine Reports. 9: 1–10.

Mullens, B. A., E.G. McDermott, A.C. Gerry. 2015. Progress and Knowledge gaps in *Culicoides* ecology and control. Veterinaria italiana. 51(4): 313–323.

Pfannenstiel, R.S., B.A. Mullens, M.G. Ruder, L. Zurek, L.W. Cohnstaedt, and D. Nayduch. 2015. Management of North American *Culicoides* biting midges: current knowledge and research needs. Vector-borne and zoonotic diseases. 15(6): 374–384.

Purse, B.V., S. Carpenter, G.J Venter, G. Bellis, and B.A. Mullens. 2015. Bionomics of temperate and tropical *Culicoides* midges: knowledge gaps and consequences for transmission of *Culicoides*-borne viruses. Annual review of entomology. 60: 373–392.

Reiskind, M. H., R. H. Griffin, M. S. Janairo, and K.A. Hopperstad. 2017. Mosquitoes of field and forest: the scale of habitat segregation in a diverse mosquito assemblage. Medical and Veterinary Entomology. 31(1): 44–54.

Relf, V. E., E.R. Morgan, J.E. Hodgkinson, and J.B. Matthews. 2012. A questionnaire study on parasite control practices on UK breeding Thoroughbred studs. Equine Veterinary Journal. 44(4): 466–471.

Rigot, T., M. Vercauteren Drubbel, J.C. Delécolle, and M. Gilbert. 2013. Farms, pastures and woodlands: the fine-scale distribution of Palearctic *Culicoides* spp. biting midges along an agro- ecological gradient. Medical and Veterinary Entomology. 27(1): 29–38.

Rivera, N.A., C. Varga, M.G. Ruder, S.J Dorak, A.L. Roca, J.E. Novakofski, and N.E. Mateus- Pinilla. 2021. Bluetongue and Epizootic Hemorrhagic Disease in the United States of America at the Wildlife–Livestock Interface. Pathogens. 10(8): 915.

Smith, K. E., and D.E. Stallknecht. 1996. *Culicoides* (Diptera: Ceratopogonidae) collected during epizootics of hemorrhagic disease among captive white-tailed deer. Journal of medical entomology. 33(3): 507–510.

Stallknecht, D.E., A.B. Allison, A.W Park, J.E. Phillips, V.H Goekjian, V.F. Nettles, and J.R. Fischer. 2015. Apparent increase of reported hemorrhagic disease in the midwestern and northeastern USA. Journal of Wildlife Diseases. 51(2): 348–361.

Swanson, D. A., and M.W. Turnbull. 2014. Molecular Identification of Bloodmeals from *Culicoides* Latreille (diptera: Ceratopogonidae) in the Southeastern USA. Proceedings of the Entomological Society of Washington. 116(3): 354–357.

Tabachnick W.J. 1996. *Culicoides variipennis* and bluetongue-virus epidemiology in the United States. Annual Review of Entomology. 41: 23–43.

Talavera, S., F. Muñoz-Muñoz, M, Verdún, N. Pujol, and N. Pagès. 2018. Revealing potential bridge vectors for BTV and SBV: a study on *Culicoides* blood feeding preferences in natural ecosystems in Spain. Medical and Veterinary Entomology. 32(1): 35–40.

Vigil, S. L., M. G. Ruder, D. Shaw, J. Wlodkowski, K. Garrett, M. Walter, and J.L. Corn. 2018. Apparent range expansion of *Culicoides* (Hoffmania) *insignis* (Diptera: Ceratopogonidae) in the southeastern United States. Journal of medical entomology. 55(4): 1043–1046.

Wegner, J. F., and G. Merriam. 1979. Movements by birds and small mammals between a wood and adjoining farmland habitats. Journal of Applied Ecology. 16: 349–357.

Wernike, K., B. Hoffmann, B., and M. Beer. 2015. Simultaneous detection of five notifiable viral diseases of cattle by single-tube multiplex real-time RT-PCR. Journal of virological methods. 217: 28–35.

Wirth, W. W., and F.S. Blanton, F. S. 1967. The North American *Culicoides* of the guttipennis group (Diptera: Ceratopogonidae). The Florida Entomologist. 50(3): 207–232.

Wirth, W. W., A.L. Dyce, B.V. Peterson, and I.T. Roper, I. T. 1985. An Atlas of wing photographs, with a summary of the numerical characters of the Neartic species of *Culicoides* (Diptera: Ceratopogonidae). Contributions American Entomological Institute. 22(4):1–46.

